# Minimally invasive electrocorticography (ECoG) recording in common marmosets

**DOI:** 10.1101/2024.10.27.620518

**Authors:** Silvia Spadacenta, Peter W. Dicke, Peter Thier

**Affiliations:** Cognitive Neurology Laboratory, Hertie Institute for Clinical Brain Research, University of Tübingen

**Keywords:** Electrocorticography (ECoG), common marmoset, Non-human primates (NHP), cortical mapping

## Abstract

Electrocorticography (ECoG) enables high spatio-temporal resolution recordings of brain activity with excellent signal quality, making it essential for pre-surgical mapping in epilepsy patients and increasingly useful in neuroscience research, including the development of brain-machine interfaces. The relatively minimal cortical convolution in non-human primate (NHP) brains facilitates ECoG recordings, in particular in the common marmoset (Callithrix jacchus), whose lissencephalic (unfolded) brain surface provides broad cortical access. One of the key advantages of ECoG recordings is the ability to study interactions between distant brain regions. The standard approach is the usage of large electrode arrays, which, however, require extended trepanations and a trade-off between size and electrode spacing. This study introduces an alternative ECoG approach for examining interactions between multiple distant cortical areas in marmosets, combining the advantages of circumscribed trepanations with high-density electrode arrays at specific sites of interest. Two adult marmosets underwent ECoG implantation across frontal, temporal, and parietal regions of interest, assessed by circumscribed trepanations to position individual high density electrode arrays. Postoperative monitoring showed rapid recovery and no long-term complications. The animals remained healthy, and stable neural recordings were successfully obtained during behavioral tasks, highlighting the method’s effectiveness for long-term cortical monitoring.

## Introduction

Electrocorticography (ECoG) is a remarkable tool for investigating neural dynamics, offering high spatio-temporal resolution insights into cortical activity. In comparison to intracortical recording, this technique is much less invasive, since the ECoG electrode arrays can be placed either epidurally or subdurally, without penetrating the cortical tissue. The recorded signal results from the summation of the electrophysiological activities of many neurons near the cortical surface.

In human medicine, electrocorticography plays a crucial role in accurately identifying epileptic foci in patients with drug-resistant epilepsy prior to resective surgery. By localizing seizure-onset zones, ECoG allows for more targeted surgical interventions, which can improve patient outcomes (Englot et al., 2012; Parvizi and Kastner, 2018). Beyond its clinical utility, ECoG recordings from these patients have also been instrumental in advancing neuroscience research, providing real-time insights into brain functions such as motor control, language, and sensory processing (Miller et al., 2010; Chang, 2015). Additionally, these recordings have been leveraged to develop brain-machine interfaces (BMIs), which aim to restore motor function and communication in individuals with paralysis or other neurological disorders (Schalk et al., 2008; Khodagholy et al., 2015; Sisterson et al., 2019), emphasizing the great usefulness of this technique.

ECoG has been also extensively used in basic research in several species, such as rats (Yoshimoto, et al., 2022), or macaque monkeys (Spyropoulos et al., 2018; Kanth and Ray, 2020). More recently, it has gained traction as a valuable tool for investigating neural dynamics in common marmosets, a New World primate species. The marmoset’s smooth (lissencephalic) brain surface makes nearly all cortical regions accessible for recording. Additionally, the marmoset’s thin dura mater supports high-quality signal acquisition through epidural ECoG arrays, eliminating the need for riskier subdural procedures that may increase the risk of cerebrospinal fluid leaks or infection (Komatsu et al., 2019; Kaneko et al., 2022).

However, it is clear that the epidural approach does not eliminate the need for a trepanation of the skull, which is always required to place the electrode array directly on the cortical surface. Given that one of the key benefits of ECoG recordings is the ability to monitor the flow of information between distant brain areas, large trepanations are often necessary. Moreover, aside from the unavoidable surgical risks or the potential introduction of germs, exposing large sections of the dura to non-biological materials may lead to undesirable reactions, such as the rapid formation of granulation tissue. This will not only soon deteriorate signal quality but potentially also affect the well-being of the experimental animal, e.g. because of an increased risk of seizures (Hauss-Wegrzyniak, et al., 2002; Cole et al. 2011; Yan et al., 2023). Hence, further refinements of the approach, which promise to reduce the likelihood of such risks and complications, remain desirable.

In this paper, we present a refinement—a method for chronic epidural ECoG implantation to study long-range interactions between cortical areas that minimizes cranial drilling while preserving the ability to document neural activity in areas of interest with high-density electrode arrays. Moreover, we propose a prophylactic antiepileptic protocol aimed at reliably preventing postsurgical seizures, which may still occur despite the approach being significantly less traumatic than traditional methods that involve extended trepanations.

## Methods

### Animals and housing

Surgeries were performed on 2 adult common marmosets (Callithrix jacchus), one male (M1, age 4 yrs and 3 months old) and one female (M2, age 4 yrs and 6 months old). The animals were housed with their respective partner in a temperature-controlled room with a 12-hour light/dark cycle. The animals were fed with a well-balanced mixture of fruits, vegetables, insects (mealworms and locusts), and arabic gum, fully satisfying their metabolic needs outside and during experimental periods. During experimental periods the food offered in the cage was gradually reduced up to 50% of the ad libitum consumption over the course of a 12 days window to increase the motivation to work for the necessary food complement provided during experimental sessions. Food availability would have been increased if a weight loss exceeding 10% of the starting weight had been observed. However, this was never the case over the course of months of training and recordings. In order to ensure that the animals were indeed able to cope with this regime, we regularly checked their behavior also outside experimental sessions in the facility. Water was available ad libitum at any time.

All experimental protocols were approved by the local animal care committee (Regierungspräsidium Tübingen, Abteilung Tierschutz) and fully complied with German law and the National Institutes of Health’s Guide for the Care and Use of Laboratory Animals.

### Stereotactic planning

Based on standard marmoset brain atlases (Paxinos et al., 2012; Bakker, et al., 2015; Majka et al., 2016; Majka et al., 2020), we determined the coordinates of the regions to be covered by ECoG electrode arrays. The chosen RAS coordinates (inter-aural (i-a), dorsal-ventral (d-v), inferior-superior (i-s)) determined the center of the 5 mm diameter electrode arrays. In both animals (M1 and M2), frontal arrays were centered between area 8 ventral (A8v) and area 6 ventral (RAS coordinates +14.5 mm i-a, +7 mm d-v, +13.4 mm i-s), while parietal arrays where centered on the lateral intraparietal area (LIP) at RAS coordinates +1 mm i-a, +6 mm d-v, +17 mm i-s). M1 received a third array, covering part of the middle temporal cortex and superior temporal sulcus (STS), centered on its fundus region (FST; Paxinos et al., 2012) at RAS coordinates +2 mm i-a, +11 mm d-v, +9.5 mm i-s.

The coordinates were projected onto a 3D skull model derived from a CT scan using CAD software (SolidWorks). A 3D-printed model was produced, with 1 mm-deep, 5 mm-diameter indentations, marking the desired arrays locations (Fig. 1A). Prior to surgery, the 3D model was mounted on a stereotaxic frame (David Kopf Instruments), which was adapted for marmosets with thinner ear bars and an elongated bite plate. Then three manipulators, each equipped with a metal rod, were mounted on the frame in order to target the desired electrode array location on the skull, orienting the rod perpendicularly to the skull at the respective target location. A schematic representation of the rods’ orientation is presented in Fig. 1B. During surgeries the manipulator configurations were reinstated. At the beginning of each array implantation, the respective rod was driven down with the manipulator approaching the skull surface. As the rods had the same diameter as the array, they allowed us to use a pencil to mark the required trepanations on the exposed bone by circling around their circumference.

**Fig. 1.**
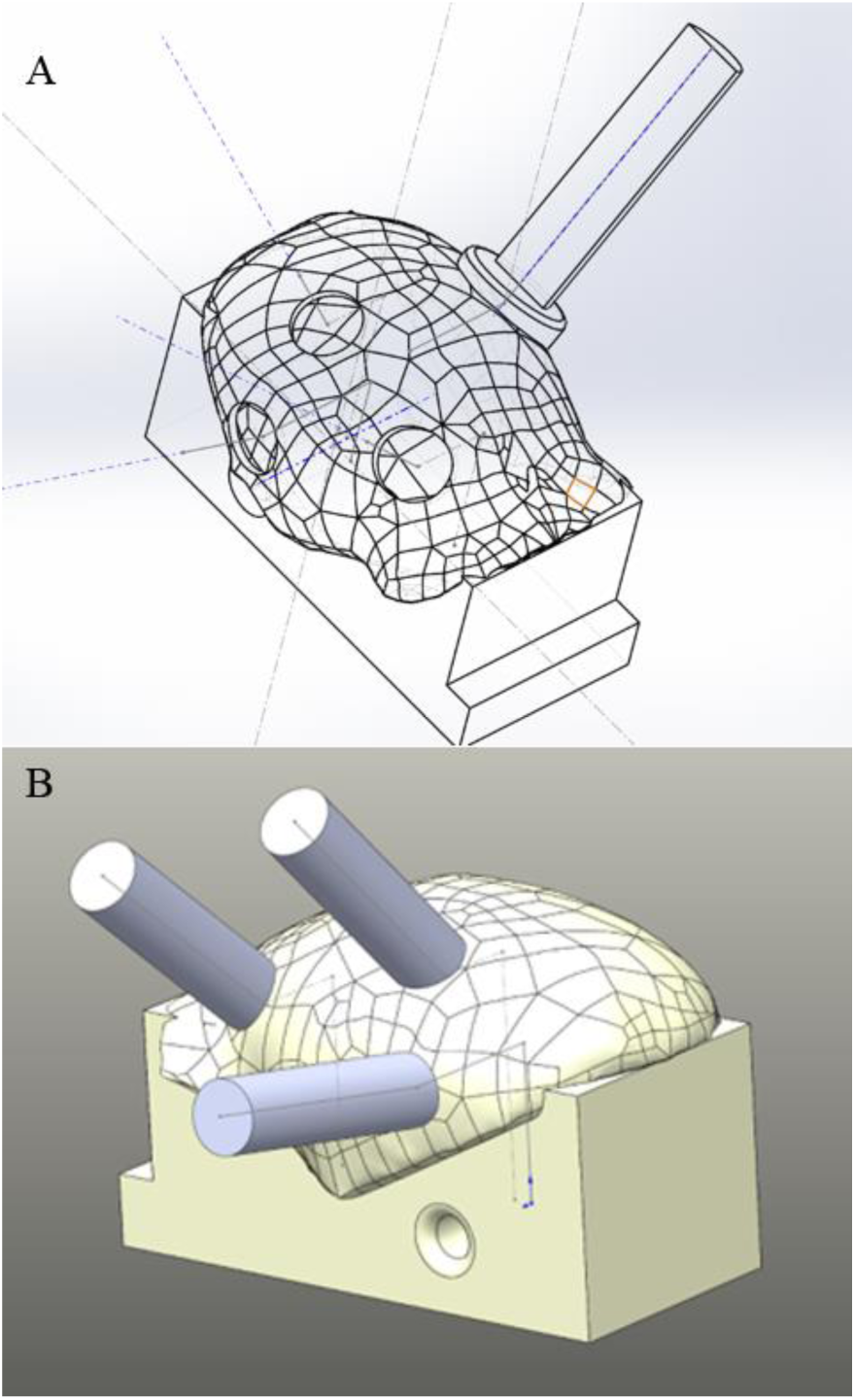
Stereotactic planning models. (A) Schematic 3D representation of a marmoset’s skull (Solid Works). The circular marks, correspond to the future trepanations for the placement of ECoG arrays and are centered on the chosen RAS coordinates. The elongated structure corresponds to a head holder, implanted in an earlier surgery to immobilize the head for behavioral training on a variety of visual and visuomotor tasks. Note that the position and orientation of the implanted head holder was chosen such as to provide sufficient space for the ECoGs arrays and their connectors. (B) Schematic 3D representation of a marmoset’s skull (Solid Works), including a simulation of the rodś positions aiming at the desired locations for trepanation during the surgery.

### Electrocorticography (ECoG) arrays

We utilized electrocorticography (ECoG) arrays manufactured by Atlas Neuroengineering (Leuven, Belgium), shown in Fig. 2A. These electrode arrays are composed of a flexible polyamide foil, 7 µm thick, designed for adaptability on the curved surfaces of small skulls, such as those of marmosets or rodents. Each comprised 32 specific electrodes, embedded in the foil in a circular arrangement. The electrodes were connected to a 32-channel Omnetics micro female socket, which offered additional plug-connections for a possible reference electrode and a ground electrode, the latter positioned independent of the foil. Each electrode had a diameter of 100 µm with the same nearest neighbor distance of 800 µm on the horizontal and the vertical axis. The total array diameter amounted to 4.8 mm. Two array configurations were used: one with an integrated reference (Fig. 2A, where the internal reference is depicted as a larger black dot outside the array) and the other with an external reference, both provided by Atlas Neuroengineering. As external reference we placed a short (1.5 cm) sterile silver wire, together with some sterile NaCl 0.9% solution, that connected the edge of the skin around the headpost with a dedicated channel. This wire was removed after each daily session. We refrained from inserting the wire chronically in order to minimize infection risks. In the first animal (M1), we used an external reference configuration, whereas in M2, ECoG arrays with an integrated reference were implanted.

**Fig. 2.**
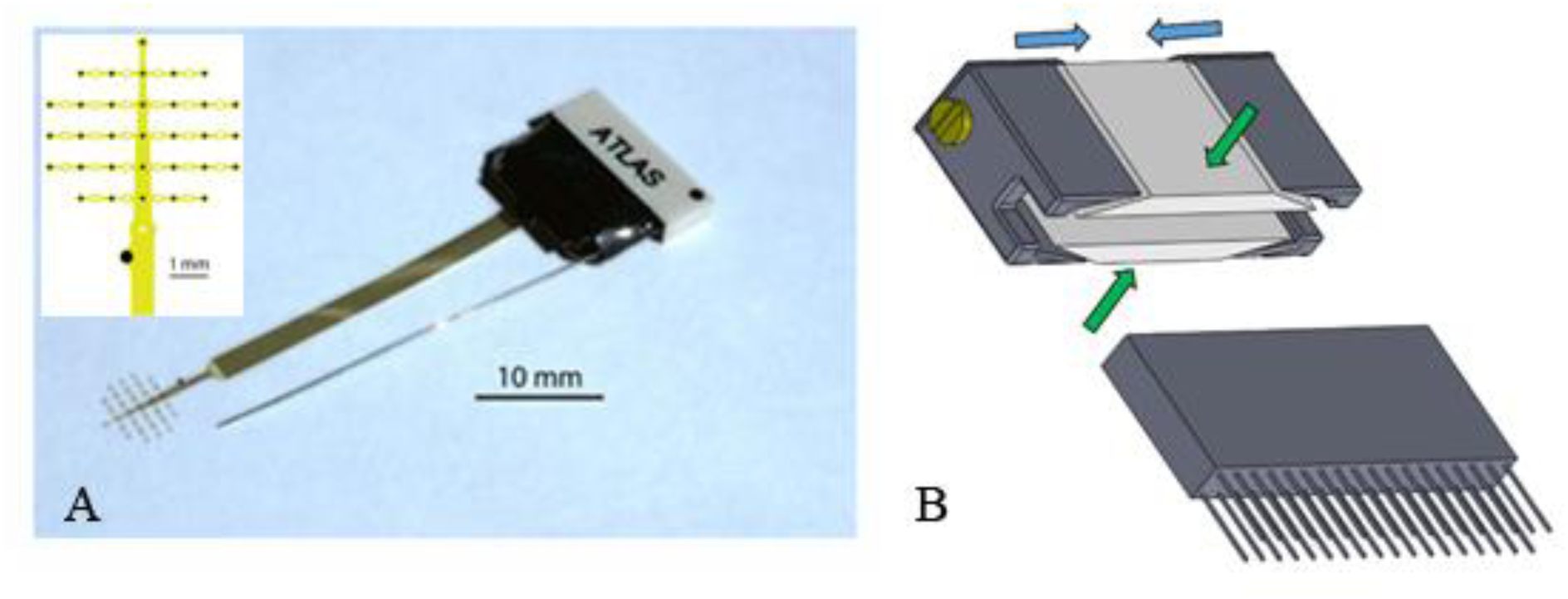
ECoG array and protection system. (A) 32 channel ECoG electrodes array with strip conductor connected to an Omnetics socket (ATLAS Neuroengineering, www.atlasneuro.com), right) and an enlarged schematic representation of the electrodes (small black dots) (left). The larger black dot on the strip conductor indicates the reference electrode. Images courtesy of ATLAS Neuroengineering. (B) Schematic representation of the cap (upper figure), protecting the Omnetics sockets outside of experimental sessions, and of the socket (lower figure). The cap features a central component made of PEEK (shown here in white) and two outer metal parts (depicted in dark grey, on the left and right). Once the cap is placed over the Omnetics socket, tightening the screw (yellow element) will bring the outer parts (dark elements) closer together (blue arrows direction), causing the two central components to approach each other (green arrows direction), thereby firmly clasping the cap to the socket. Conversely, loosening the screw disengages the clamping mechanism, allowing the protective cap to be easily removed.

To protect the sockets when not in use, they were covered with a customized cap made of plastic (PEEK) and aluminum. To ensure that the cap would not be removed by the animal, a small screw was added to it. The screw allows the two side components (see Fig. 2B upper part), dark grey cap elements) to be brought closer together, causing the two central components to approach each other, ensuring that the cap is firmly fastened to the Omnetics socket.

## Surgery

### Anesthesia Management

To prepare the animals for anesthesia, access to food was limited to a small amount of moist food, such as grapes, a few hours before surgery. Water was provided ad libitum. After transfer to the surgery room in their familiar transport box, anesthesia was induced with sevoflurane (2.5 – 8% in > 90% oxygen) allowed to flow into the box. This approach ensured a stress-free induction as the animals were accustomed to the transport box from their daily training. Pre-medication by injection was deliberately avoided to reduce the stress associated with handling the animal. Once anesthesia was induced, the animals were taken out of the box, and anesthesia was maintained with a well-fitted face mask delivering sevoflurane in oxygen. To intensify analgesia and muscle relaxation, intramuscular ketamine (15 mg/kg) and midazolam (0.1–0.2 mg/kg) were administered, with their dosage chosen based on the depth of anesthesia. If anesthesia was sufficient, intubation followed with a silicon tube (diameter 2–2.5 mm) whose tip was cut slanted. This custom-made endotracheal tube was inserted through the larynx under visual guidance. To prevent irritation and consecutive larynx closure when trying to insert the tube, the local anesthetic lidocaine (1%) was sprayed on the larynx and surrounding tissue. Once the tube was in place, it was connected to the ventilator, and the animals were ventilated with sevoflurane (2.5–5%) in a mix of oxygen and air (3:1 to 2:1). Throughout the procedure and the subsequent surgery, respiration, heart rate and blood pressure were continuously monitored. In order to maintain proper hydration lactated Ringer’s solution was administered intravenously via the femoral or tail vein at an initial rate of 3–5 ml/kg/h and later adjusted based on the animal’s needs, guided by blood pressure and other physiological indicators. Once a stable i.v. line was available, anesthesia was switched to a combination of sevoflurane (2–3%) in 50% air and oxygen, alongside remifentanil (50 µg/kg/min) and propofol (5–10 mg/kg/h). Body temperature, heart rate, and exhaled CO₂ levels were closely monitored and parameters adjusted as needed. Postoperative medication comprised the NSAID meloxicam (0.2 mg/kg), dexamethasone (0.5–1 mg/kg), and the antiemetic maropitant (1 mg/kg), along with broad band antibiotics administered for 5 days. After surgery, continuous monitoring was carried out both in person and by camera for several hours. Following this, the monkeýs condition was assessed regularly multiple times during the day until 18:30 (lights off in the animal facility) for at least 10 days to ensure early detection of any complications or discomfort of the animal.

### Headpost implantation

As mentioned earlier, several months before the implantation of the ECoG array, the marmosets were fitted with a headpost on the side contralateral to the planned recording sites. This setup was necessary to enable painless fixation of the head, which was required for the visual and visuomotor tasks the animals needed to master. The titanium headpost was fixed to the skull with three bone anchors (Fig. 3A) and resin cement (3M ESPE, RelyX^TM^ Unicem 2 Automix, Self-Adhesive Resin Cement) linking the headpost, the screws and the bone. First, three 1 x 2 mm slits were cut into the skull with an ultrasonic cutting tool (Mectron, Piezosurgery). The slits enabled us to insert the flat bar (3 x 1.5 x 0.5 mm) of an elongated T-shaped anchor parallel to the slit into the epidural space beneath it (Fig. 3B). By rotating the vertical shaft of the anchor (metric 1.4 mm threat, 4 mm length) by 90°, the anchor bar was locked securely under the bone (see Fig. 3C, showing the anchor plate position relative to the bone after 90° rotation) with a nut which gently applied pressure to the bone from above (Fig. 3D). Once all three anchors were in place, the bone in between the anchors and next to them was covered with a layer of Super-Bond (SUN Medical) following meticulous cleaning of the area. The Super-Bond layer served as the foundation for a UV-cured resin bed (3M ESPE, RelyX^TM^ Unicem 2 Automix, Self-Adhesive Resin Cement) that was spread over it to embrace the headpost basis positioned in between the anchors as well as the vertical anchor shafts. Finally, the surrounding skin was repositioned such as to cover most of the cement.

**Fig. 3.**
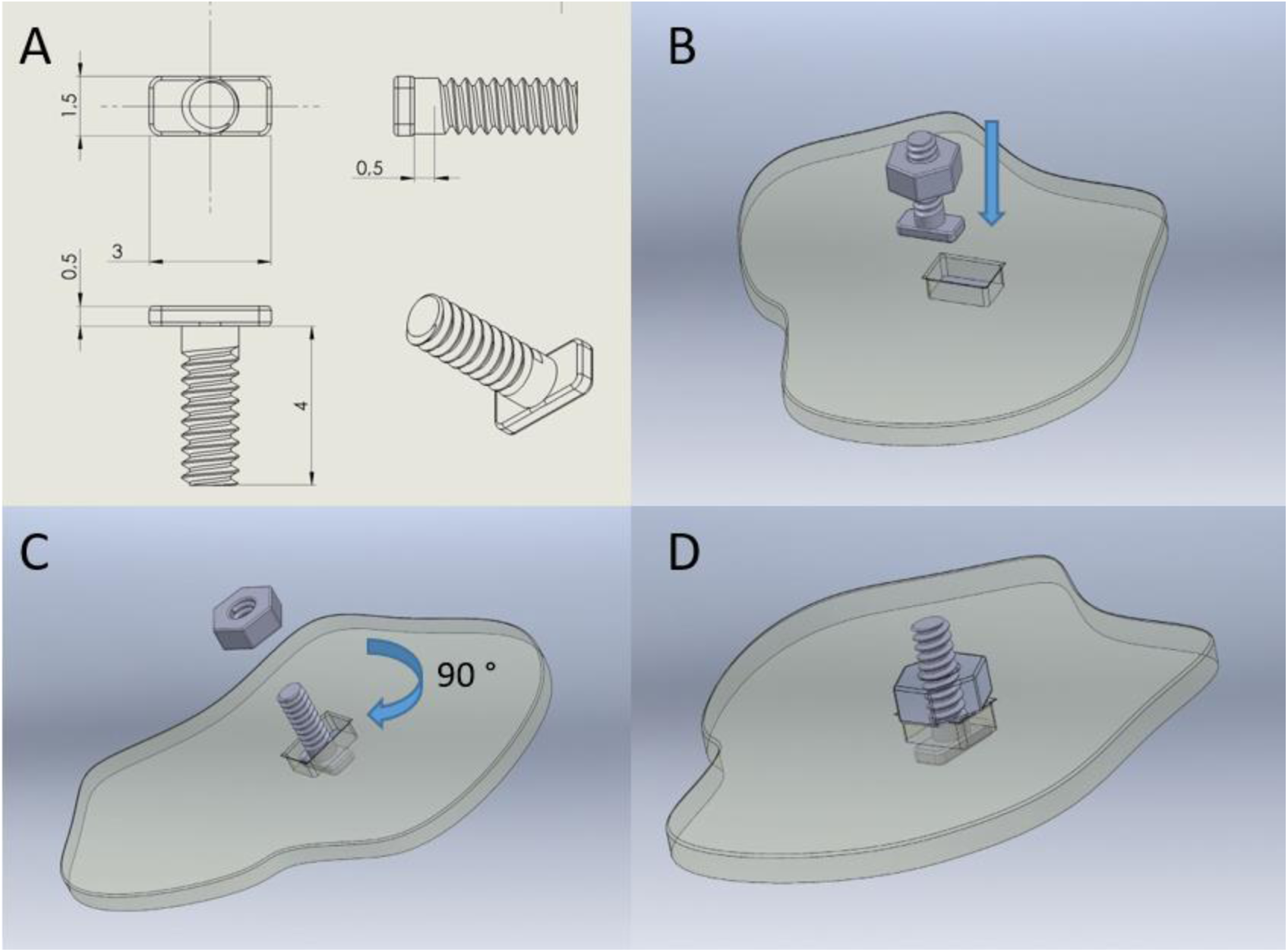
Design and insertion procedure of a custom titanium bone anchor. (A) Drawing of the titanium (Ti) anchor, cut from a (metric) 1.4 mm screw. The head was partially removed to shape a thin and flat footplate. (B) Anchor together with a Ti 1.4 mm metric nut over a made-to-measure bone slit needed to insert the footplate under the skull. (C) After inserting the footplate, the anchor was turned by 90 degrees and (D) fixed to the skull by gently tightening the nut.

### ECoG array implantation

As first step, the marmosets were positioned in a stereotaxic frame (Kopf, USA). For each array we adopted the following procedure. After opening the skin overlying the calotte, the target location was identified using the pre-set stereotactic rods and manipulators and the boundaries of the trepanation delineated as described before (see Stereotactic planning section). The trepanation was performed using a manually held ultrasonic knife which minimizes the risk of inadvertent damage to the underlying dura. The resulting circular bone flap was transferred into cold and sterile saline 0.9% for later repositioning. Following careful cleaning of the exposed dura from residual bone debris, the ECoG electrode array was gently placed on the dura with the output cable oriented toward the headpost. The cut-out bone flap was then repositioned and stabilized relative to the surrounding bone with the aid of a thin, V-shaped titanium wire of 0.5 mm diameter (Fig. 4A). The vertex of the V-shaped wire was positioned on the center of the bone flap while the two ends of the wire were placed over the surrounding bone. The wire was then firmly connected to the flap and the neighboring bone by applying small drops of RelyX^TM^ resin cement on the wire vertex and the two ends. This procedure was repeated for each ECoG array, starting with the parietal location, followed by the frontal and finally the temporal location, the sequence chosen to accommodate the different distances in view of a fixed cable length.

**Fig. 4.**
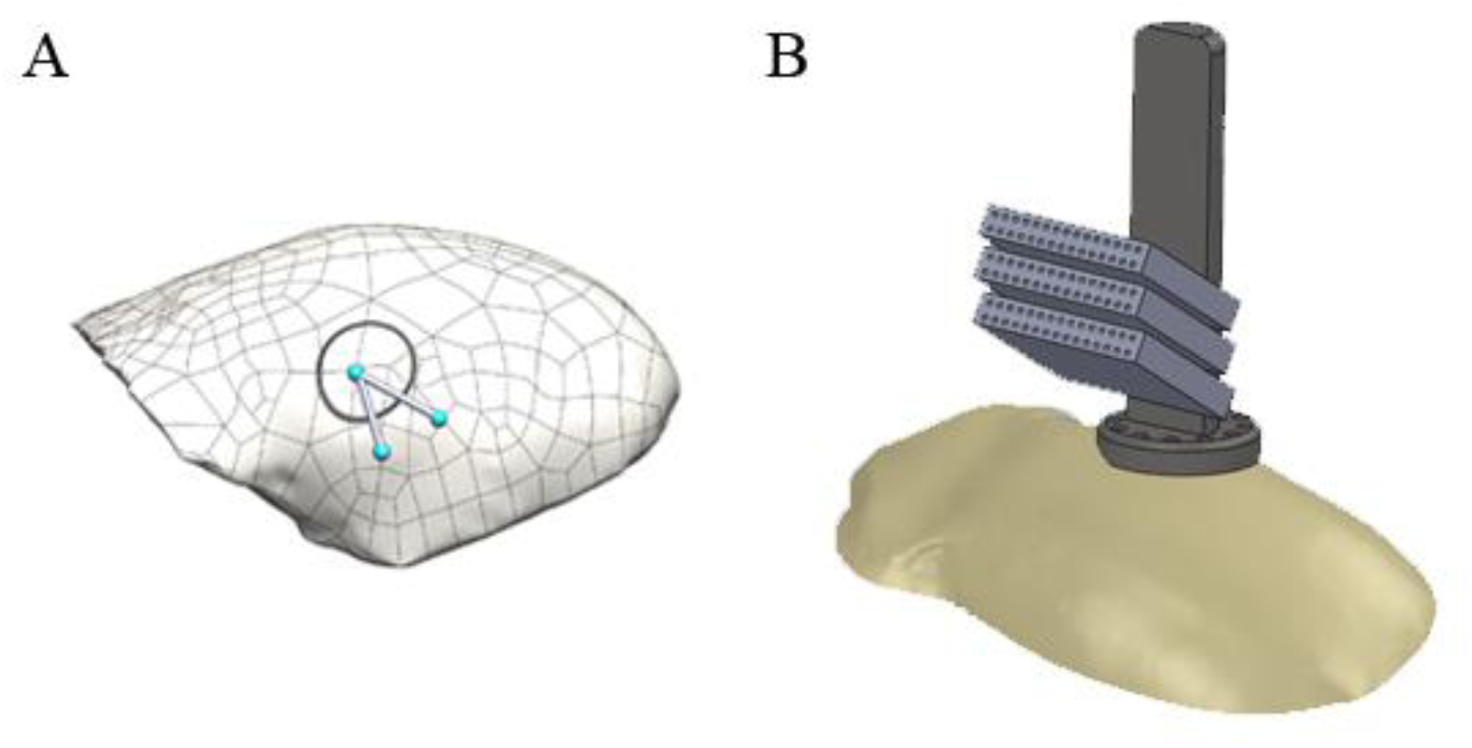
Bone flap stabilization and sockets positioning. (A) Schematic representation of the V-shaped titanium wire used to secure a bone flap reponed after the placement of the ECoGs such as to prevent the flap’s descent onto underlying cortex. The blue dots indicate the points where the wire was attached to the bone using resin cement. Care was taken to ensure that the titanium wire was oriented such as to not overlap with the ECoG wire path leading to the headpost. (B) Schematic representation of the stack of ECoG array sockets fixed to the headpost with RelyX^TM^ resin cement (not shown). From top to bottom: parietal, frontal, and temporal sockets.

The sockets were attached to the headpost with RelyX™ resin cement on top of each other (Fig. 4B), with the parietal array socket positioned highest (farthest from the headpost base), followed by the frontal socket, and the temporal socket positioned lowest. Importantly, the sockets were spaced adequately to accommodate all the corresponding headstages during recordings. In order to accommodate excess cable lengths in the case of the two arrays that were placed relatively close to the headpost (the frontal and even more so the parietal array), their cables were folded in several layers close to the headpost and embedded in cement. Crucially, once a cable was bent in a particular direction, this orientation was consistently maintained and never reversed, as doing so could potentially damage the cable’s structure. If a cable did not remain securely attached to the bone as it was routed toward the headpost, we applied a small drop of RelyX™ resin cement to bond it to the bone. After having positioned all arrays and secured the cables and sockets, we repositioned the skin and sutured it.

### Postoperative care

The animals were closely monitored for any signs of discomfort or pain, both in person by an experimenter and through continuous 24-hour video surveillance starting immediately after surgery. Approximately 30 minutes after having awoken from anesthesia, M1 experienced a tonic-clonic seizure affecting the left arm and leg (opposite to the side of the ECoG implantations). The seizure lasted about 2 minutes and was promptly treated with 0.0125 mg / 100 gr of diazepam. The episode quickly subsided, followed by postictal drowsiness. Three days post-surgery, M1 experienced again a sequence of focal tonic-clonic seizures of milder intensity as compared to the first post-surgery event (left visual hemineglect and left arm/leg paresis). Each bout lasted for a couple of seconds to a minute and was followed by postictal drowsiness. These seizure bouts occurred at intervals of 3-4 hours, during which the animal exhibited normal behavior as, for instance, demonstrated by the ability to feed independently, to accept treats, and to interact as usual with its cage mate. The seizures were observed only during the morning hours and early afternoon.

While initial attempts to stop seizures with diazepam at the same dosage that resolved the post-surgery seizure (0.0125 mg / 100 g) may have shortened their duration, this drug failed to prevent new seizure events and seemed to worsen postictal drowsiness. This is why we transitioned to Levetiracetam (LEV), an antiepileptic drug, which has been shown to reduce L-DOPA-induced dyskinesia in MPTP-lesioned marmosets (Hill, et al., 2004). We began with a dosage of 1 mg of LEV per 100 g (100 mg/kg, 0.01 ml/100g), administered orally twice a day, which is equivalent to the minimum pediatric dose in human medicine. The drug was given either with marshmallow/grape juice or diluted in arabic gum. Within 24 hours of the first dose, seizures ceased, and the animal appeared unremarkable in every respect thereafter. Over the following six weeks, the dosage was gradually reduced by half each week without the seizures returning. Twenty months post-surgery, M1 remains completely healthy in every respect.

To prevent a similar sequence of full-fledged seizures, we decided to subject M2 to a prophylactic anticonvulsant treatment with LEV for 4 weeks, after observing what might have been a single subliminal focal seizure on day one post-surgery. Since then, M2 has remained healthy and seizure-free for over a year. Neither M1 nor M2 showed any drug-related side effects and exhibited a completely unremarkable behavior also during the weeks of medication.

### Data Acquisition and experimental setup

Simultaneous ECoG recordings were obtained from three brain areas in M1 (frontal, parietal, temporal) and one (frontal) in M2 respectively by using the Open Ephys data acquisition system for sampling and storing the data at 3kHz. Although also M2 had 2 arrays, unfortunately, the second array could not be accessed as its socket got destroyed by the animaĺs mate soon after surgery (see further). The headstages (RHD 32-Channel Recording Headstages, Intan technologies US) of the Open Ephys were directly connected to the Omnetics sockets while the animals were sitting in their chair with their head being immobilized. We shielded the headstages to reduce the capture of noise, in particular 50 Hz interference from the power grid. Both animals, already accustomed to perform behavioral tasks while head-fixed over the months preceding the ECoG implantations, adapted quickly (in less than a week) to the additional procedure of connecting the headstages to the sockets and their behavioral performance remained at pre-surgery levels. Head-fixing and connecting the headstages took approximately three minutes, including the time to place the external reference for M1’s ECoG arrays, which did not have an integrated reference contact (for details, see the Electrocorticography (ECoG) array section).

The recordings were performed in a small sound proof room in daily sessions lasting between 30 min and 1 hour, while the animals were sitting in a comfortable monkey chair, placed on a table facing a small computer screen (Beetronics, 10 Inch Monitor, 220 × 134 mm, 1920 × 1080 Hz resolution, framerate 60 Hz), at a distance of 26 cm. The experimental stimuli were presented with a custom made program (Nrec). Eye movements were tracked using the ISCAN ETL-200 eye tracking system, with a camera positioned on the right side of the screen, and the data were resampled at 1 kHz. Reward (either marshmallow juice, mixed with a small amount of arabic gum and banana powder or arabic gum diluted in water) was delivered via a small cannula placed in front of the animals’ mouths, either on or very close to the upper lip, depending on their preference. The reward delivery was controlled via a syringe pump.

ECoG signals were processed using custom MATLAB scripts and additionally the open-source software EEGLab (https://sccn.ucsd.edu/eeglab/index.php) for preliminary screening. Raw data presented here were first filtered in two stages: first, high-pass filtered at 0.1 Hz using a 4th-order Butterworth filter to remove DC, followed by a 6th-order Butterworth filter low-pass filtered with 250 Hz cut-off frequency. Residual 50 Hz power line noise and its harmonics were eliminated using an IIR comb notch filter (2.5 Hz bandwidth) in MATLAB.

## Results

All ECoG arrays functioned well and provided high-quality data for 20 months without noticeable degradation. Example of raw traces (filtered between 1 and 150 Hz for presentation purposes) are shown in Suppl. Fig. 1 (M1 parietal array, Ch1–Ch32, and temporal array, Ch33–Ch64) and Suppl. Fig. 2 (frontal arrays, M1, Ch1-Ch32 and M2, Ch33-Ch64). After approximately 8 months from implantation, four channels of the temporal array of M1 started to exhibit increased noise; however, gently repositioning the headstage in the socket often restored a good signal-to-noise ratio, indicating that this problem most likely was due to an unstable plug connection. Eventually, the original signal-to-noise ratio of these channels could no longer be fully restored. Yet, they still delivered usable signals after appropriate filtering.

After nine months of successful recording, we were no longer able to record from the frontal array in M1 due to a sudden break in its socket while the animal was freely moving in the cage. Based on our experience the transition between the white and black part of the socket (see Fig. 2A) can be fragile and may lose stability after plugging the headstages in and out for many recording sessions. We therefore highly recommend to stabilize the sockets in generous resin cement pouches.

In M2, we were only able to record data from the frontal array because the animal’s cage mate tampered with the implant by removing the initial protective cap, which lacked a screw, and chewed on the socket of the parietal array. This incident prompted us to develop the cap design depicted in Fig. 2B, which not only protects the cap from dirt but also prevents removal by other animals in the cage. As there was no way to connect a new socket to the probably still intact parietal ECoG array, we attempted to implant a new array. This allowed us to realize that the gap between the bone flap and the calotte had meanwhile partially closed by ossification. Upon opening, we observed that the old array was securely adhered to the dura. Anticipating that attempts to remove it could potentially damage the dura, we opted not to replace the array. Instead, we chose to reposition the bone flap and suture the skin closed.

To functionally characterize the areas covered by the ECoG arrays, brain activity was recorded while the animals performed various tasks. One task involved a 1-second inter-trial period followed by fixation on a central dot (0.2° visual angle) for up to 500 ms, after which visual stimuli from different categories (faces, scrambled faces, objects, body parts; size 12 x 12°) were randomly presented for durations of 300, 500, or 700 ms. Clear responses to the presentation of these visual stimuli were registered in all arrays and channels. Fig. 5 exemplifies the averaged evoked related potentials (ERPs), relative to stimulus onset for an exemplary fixation session (stimulus duration 300 ms, all stimulus categories pooled: n= 157 trials; black dashed line: fixation dot on; green dashed line: stimulus on; magenta dashed line: stimulus off). In both monkeys, the amplitude of signal changes in response to stimuli was significantly smaller in the frontal arrays compared to the parietal and temporal arrays. The consistency of the data collected from both monkeys indicates that the significantly reduced ERPs amplitudes are a characteristic of visual stimulus processing in the frontal cortex rather than a result of technical artifacts.

**Fig. 5.**
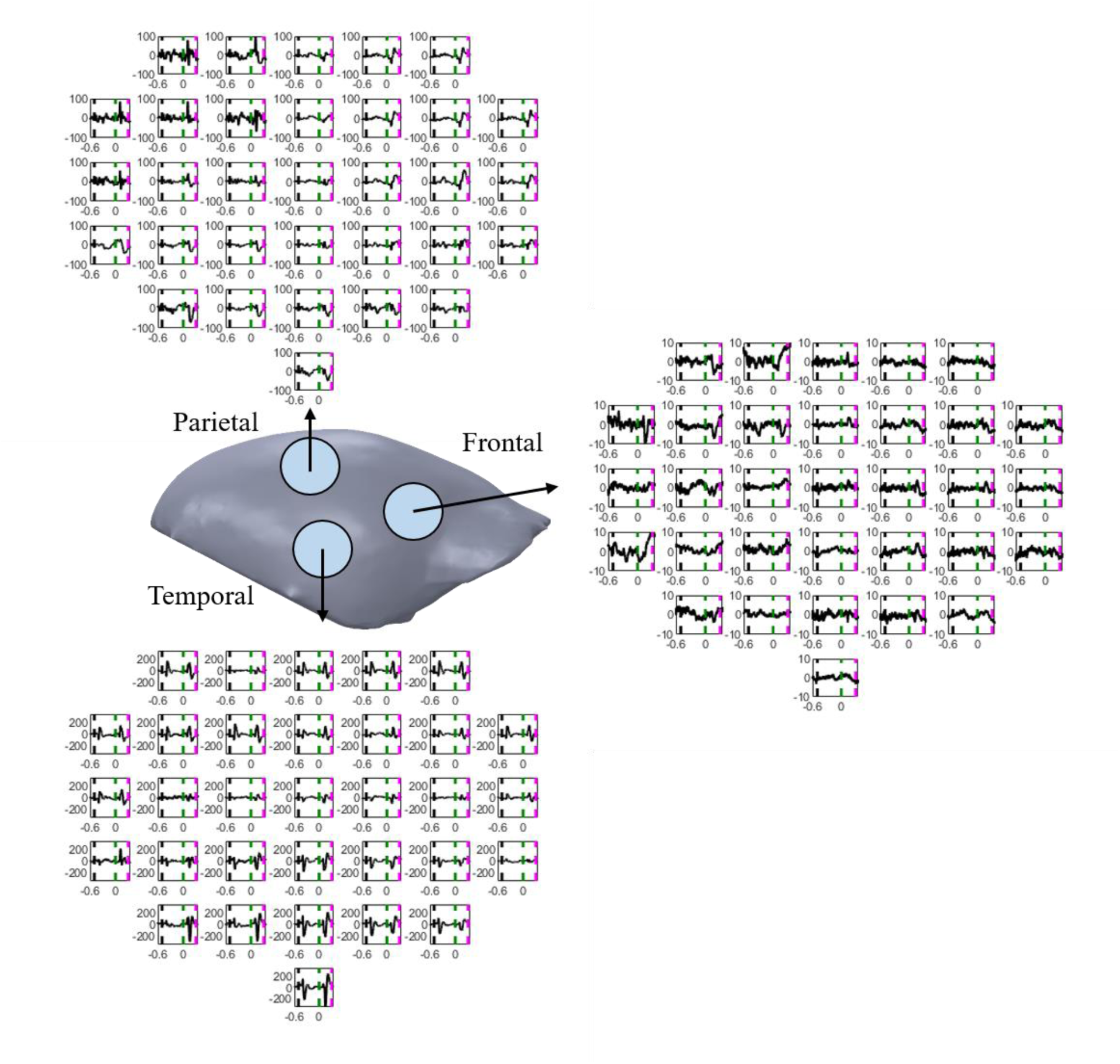
Examples of ERPs recorded from the three ECoG arrays (parietal, temporal, frontal) in M1, aligned to the onset of visual stimuli from different categories (faces, scrambled faces, objects, body parts; black dashed line: fixation dot on; green dashed line: stimulus on; magenta dashed line: stimulus off). One exemplary session is shown, with responses to various stimulus categories pooled (total number of trials n = 157; stimulus duration 300 ms). The ERPs are arranged according to the positions of their respective electrodes, and the circles on the calotte are indicative of the three arrays’ positions.

We also explored the responses to visual stimuli in the time-frequency domain using a continuous Morlet wavelet transform within the 2–150 Hz range (Cohen, 2019), with the number of wavelet cycles increasing with frequency (from 3 to 45) to balance temporal and spectral resolution. This analysis produced a time-frequency representation, from which power was extracted and baseline-corrected to assess stimulus-related neural activity (Fig. 6). Visual stimulus onset elicited distinct time and frequency responses depending on the electrode location. Activity evoked by stimulus onset was primarily found in the theta and beta bands, while high-frequency oscillations in the gamma range (up to 150 Hz) were prominently observed across multiple channels. Notably, this high-frequency activity indicates that the signals detected by the epidural electrodes were not significantly diminished by the intervening dura mater. The fact that these activity patterns stayed stable over many months also indicates minimal or absent growth of granulation tissue between electrodes and cortex. The clear association of evoked activity involving the gamma and high gamma bands at a latency that was too short to reflect EMG activity driving movements or increased muscle tension, speaks against a relevant contamination of the signal during fixation.

**Fig. 6.**
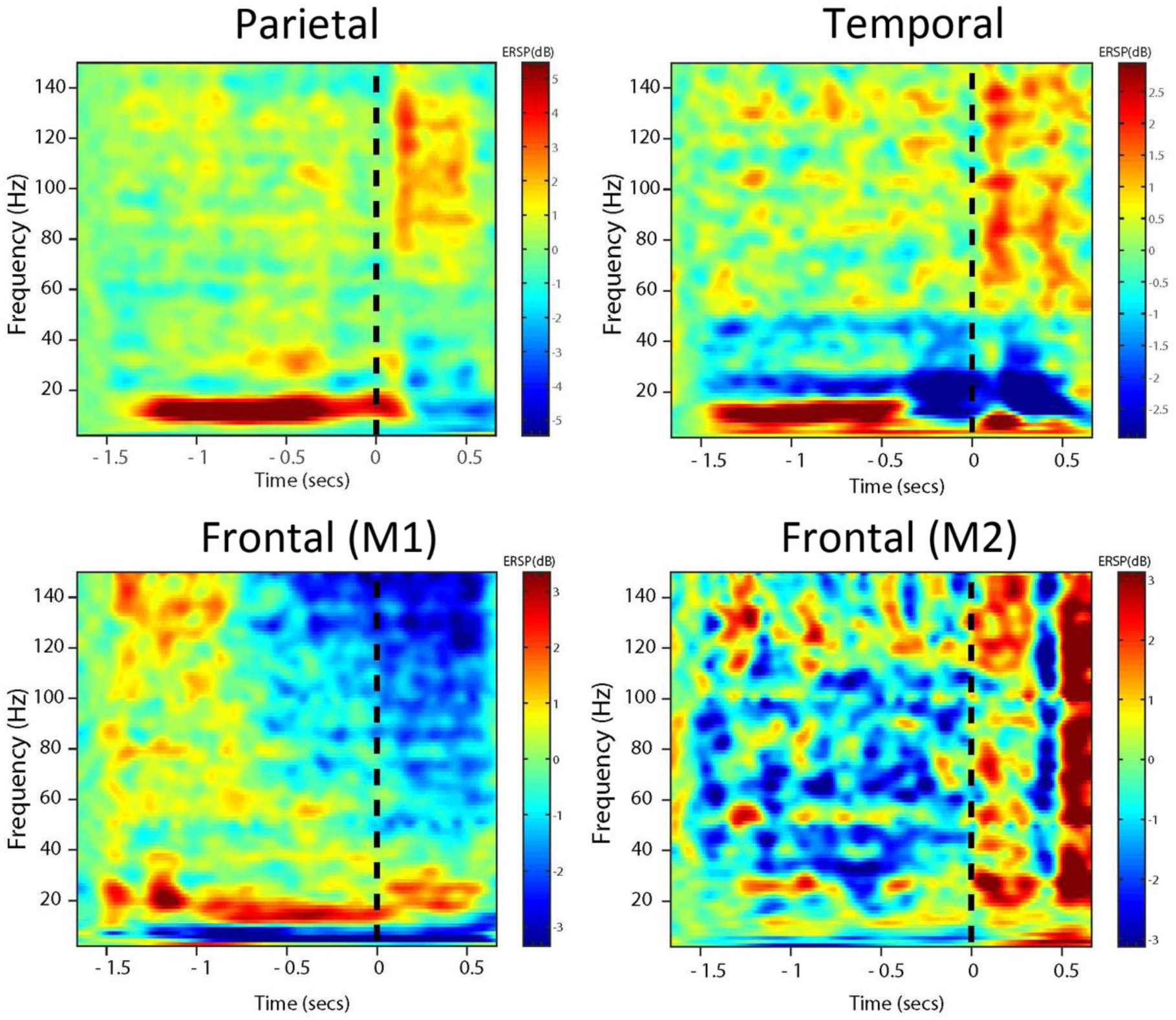
Time frequency response (2 – 150 Hz) of a representative channel for each ECoG array of M1 (parietal, temporal, frontal) and the frontal array of M2 based on an exemplary session (data pooled across stimulus categories, n = 157 trial, stimulus duration 300 ms) in which the monkeys were exposed to presentations of visual stimuli from various categories (faces, scrambled faces, objects, body parts; size 12 x 12 degs). Plots are aligned to visual stimulus onset.

On the other hand, during reward release, a high-frequency broadband signal (100–250 Hz) was observed in several frontal and parietal channels in M1. As the temporal array that was much closer to well-developed masticatory muscles (M. temporalis and masseter) did not show comparable high frequency activity, we suspect that the reward related high frequency burst was probably not a reflection of EMG activity from these muscles. Rather, we suspect an electrical interference caused by the contact between the tongue and the reward delivery spout. In contrast, M2’ s frontal array showed no reward-related artifacts, likely due to the internal reference reducing external noise contamination. However, given that EMG frequencies overlap with a broad range of frequencies, including the gamma band (Muthukumaraswamy, et al., 2013), EMG contamination cannot fully be ruled out. However, these artifacts did not interfere with the analysis, as they fell outside the relevant time windows of interest. Higher-amplitude broadband noise in the same frequency range was generated by the animals’ movements toward the end of the task, likely due to substantial EMG contamination. At this stage, the animals often lose interest in the tasks and frequently attempt to reach the reward spout with a hand, accompanied by shoulder and ear pinnae movements, clearly signaling their intent to stop. These movement-related artifacts, affecting the very last trials of a daily session, were always excluded from further analysis. Unlike the reward-related noise, these artifacts were more prominent on temporal electrodes, further suggesting the EMG nature of these broad band noise. Accompanying EMG recordings from masticatory and shoulder muscles might be considered in the future to increase confidence in the interpretation of gamma band activity as suggested by Muthukumaraswamy et al. (2013).

## Discussion

The present study introduces a novel method for chronic epidural electrocorticography (ECoG) in common marmosets, aiming to refine existing techniques in an attempt to reduce the probability of complications and to expand the duration of unrestricted signal acquisition. ECoG recordings are typically used to study the information flow between distant cortical regions. To this end, usually a single large ECoG array is placed onto cortex, requiring a large trepanation. Such large trepanations not only entail longer periods of full postsurgical recovery but also significant risks of complications due to bone necrosis, infections, chronic non-inflammatory reactions as a result of insufficient biocompatibility, and potentially the promotion of focal seizures. Considering these risks is particularly important for marmosets, as stress and postoperative recovery are significant concerns due to their sensitive nature. Our approach reduces these risks to a considerable degree by placing high density ECoG arrays epidurally over regions of interest, accessed by small, individual trepanations. To prevent the reinserted bone flap from caving in and causing pressure, we used titanium wires to stabilize it at the same level as the surrounding bone. The wire was secured to the flap and the adjacent bone using a dot-applied, acrylic-based adhesive, and the area was subsequently covered with the original muscle and skin. Our admittedly limited experience with 5 trepanations that were sealed in this way suggests that this procedure is well tolerated without any complications. Moreover, the one trepanation that was revisited after 6 months in order to replace an array showed that new bone had already closed most of the gap between the bone flap and neighboring bone. The absence of aversive effects on the underlying brain due to pressure or inflammation is well-documented by our ability to record rich patterns of local field potential activity, right after 10 days of postsurgical recovery without any noticeable changes over many months of recordings from all successfully implanted arrays. Only 4 channels of M1 temporal array showed an increased noise after 8 months of recordings, but for reasons given in the results, we would attribute the decrease in this case to the connector rather than the array.

The observation of overt focal seizures in one animal, immediately after surgery, suggests that already the minimal and fully reversible impact of the surgical manipulations needed may be sufficient to reduce the threshold for seizures, at least temporarily. While we were unable to relate the semiology of the epileptic fits to specific trepanations, their lateralization leaves little doubt that at least one of them may have been causal. In any case, anticonvulsant treatment with LEV turned out to be very effective in preventing further seizures without entailing any noticeable side effects. Hence, in order to prevent postsurgical seizures in the first place, we strongly recommend to adopt a prophylactic administration of LEV in the first weeks after surgery. This recommendation follows standard approaches in human neurosurgery (Gokhale, et al., 2013; Greenhalgh, et al., 2020). The incidence of post-craniotomy seizures in humans is highly variable, making individual risk difficult to predict. Most neurosurgeons recommend prophylactic anticonvulsive medication (Dewan et al., 2017; Spena et al., 2017), and, currently, LEV is preferred due to its favorable pharmacokinetics, tolerability, safety, and interaction profile compared to older AEDs such as phenytoin, carbamazepine, or valproate (Konrath et al., 2022). Although we did not take this prophylaxis up right after surgery in M2, it was started early enough to prevent the emergence of full-fledged focal seizures in our second animal.

Independent of being less complication prone, implanting several spatially separate ECoG arrays has the advantage of allowing the concentration of electrodes in regions of interest. We deployed arrays with a diameter of 4.8 mm comprising of 32 individual electrodes with a site of 100 µm and an inter-electrode distance of 0.8 mm. The small electrode size and short distance between electrodes ensures optimal spatial resolution in the region of interest, combined with the ability to capture the communication with other, spatially separated regions. Implantation of large ECoG arrays comes at the cost of losing spatial resolution, as these arrays usually have a much wider inter-electrode distance, typically in the range of 2 to 5 mm or higher (Slutzky, et al., 2010; Nagasaka, et al., 2011; Wang, et al., 2016; Komatsu, et al., 2019; Kaneko, et al., 2022). In our case, the only reason not to use even denser electrode arrays or to implant more than three was our initial fear that the limited space available for the omnetics sockets needed to tap the electrode signals might not suffice. However, further additions to the stack of connectors may be feasible in the future, as the animals readily integrated the extension of their head due to the omnetics mount and the neighboring headpost into their body schema.

Crucially, reduced surgical trauma and a lower risk of complications benefit animal welfare, which is essential for longitudinal studies involving behaving animals, where the health and well-being of the subjects over extended periods is critical. We highlight that, after significant time post-implantation (20 months for M1 and 15 months for M2), the ECoG arrays continue to produce signals of consistent quality, similar to those recorded in the initial days. Both animals remain in excellent health. This sustained signal quality indicates that chronic sterile inflammation and reactive granulation tissue formation, which could compromise signal integrity, have been minimal or absent. The valuable experience gained from five array implantations has allowed us to optimize the surgical procedure, ensuring minimal trauma, stable long-term signal quality, and effective seizure prevention. We believe that these refinements will not only benefit research in marmosets but could also be adapted for use in other species, thereby broadening the applicability of ECoG in neuroscience research. Future studies should continue to explore further optimizations and applications of this method, particularly in the context of brain-machine interface development and the study of complex neural networks.

## Authors’ contributions

S.S., P.W.D., and P.T. developed the method, performed the surgeries, and contributed to writing the paper. S.S. trained the animals, collected, and analyzed the data.

## Acknowledgments

We would like to extend our gratitude to Prof. Chen Chih-Yang, Kyoto University, Japan, for his invaluable discussions and insightful feedback throughout the development of our ECoG technique and to the veterinary team for their essential support in caring for the animals throughout the study.

