## Supplementary figures and images for "Minimally invasive electrocorticography (ECoG) recording in common marmosets"

### Supplementary fig. 1

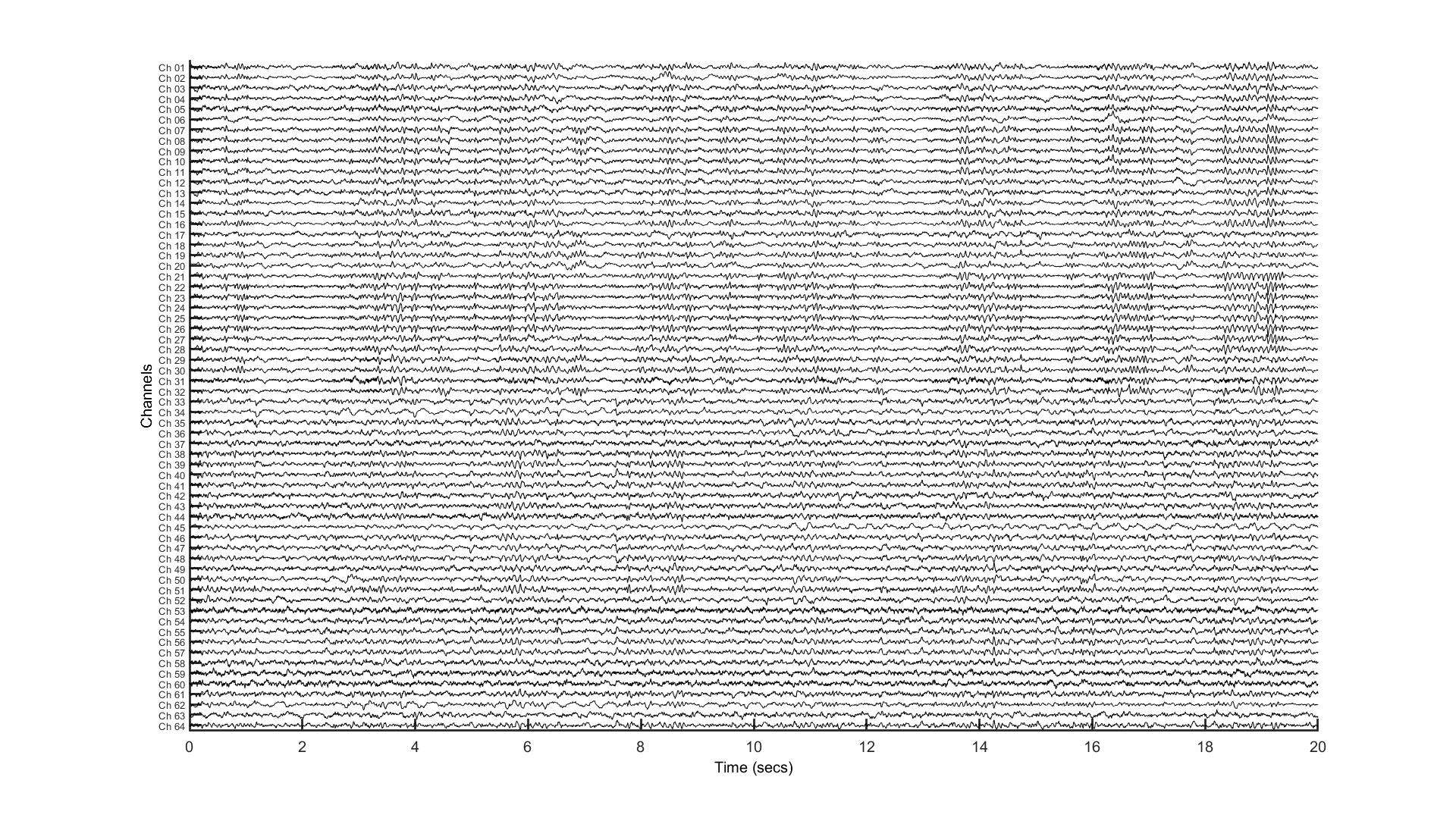

### Supplementary fig. 2

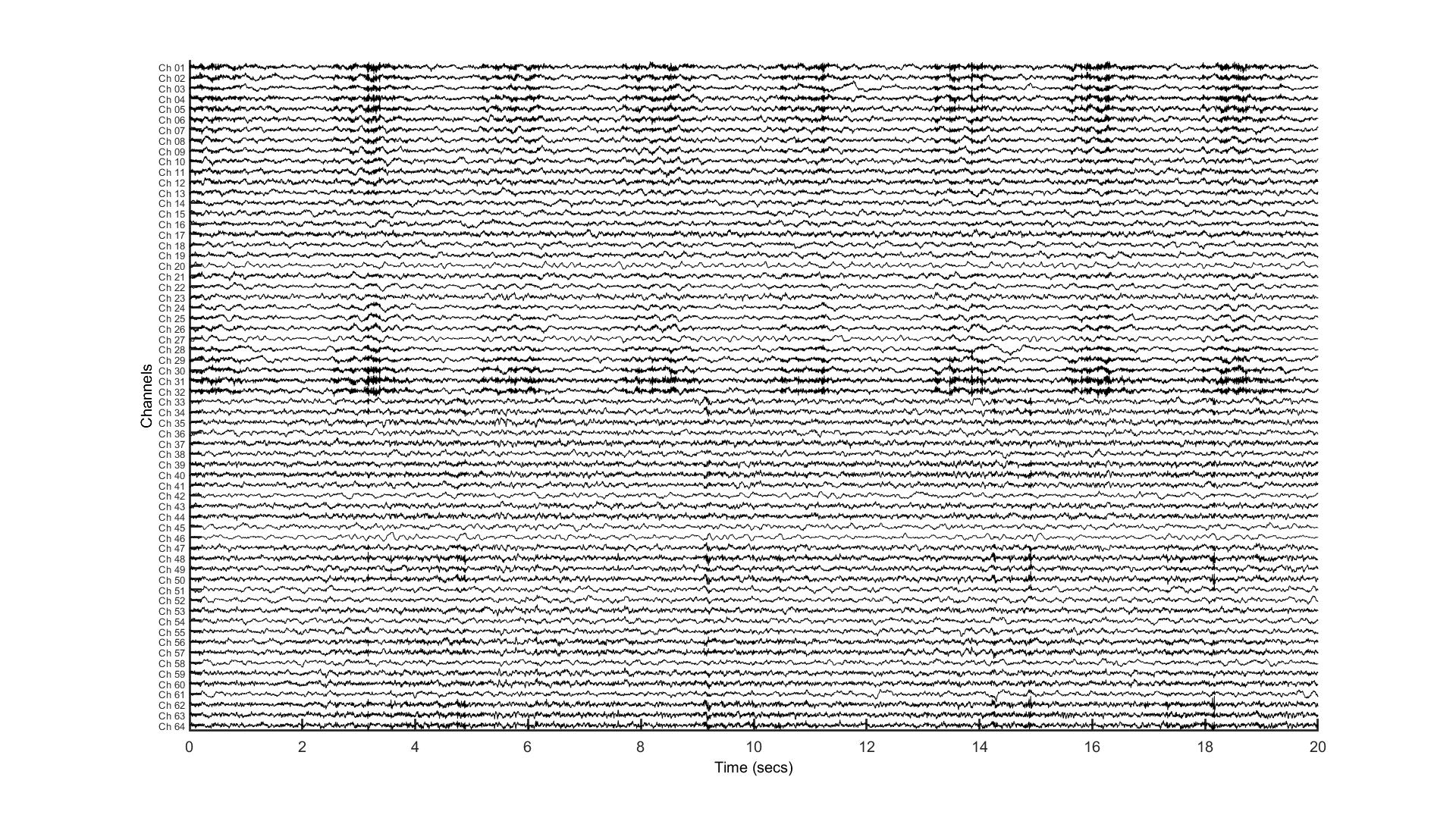
